# Robust Estimates of Overall Immune-Repertoire Diversity from High-Throughput Measurements on Samples

**DOI:** 10.1101/024612

**Authors:** Joseph Kaplinsky, Ramy Arnaout

## Abstract

The diversity of a person’s B- and T-cell repertoires is both clinically important and a key measure of immunological complexity. However, diversity is hard to estimate by current methods, due to inherent uncertainty in the number of B- and T-cell clones that will be missing from a blood or tissue sample by chance (the missing-species problem), inevitable sampling bias, and experimental noise. To solve this problem we developed Recon, a modified maximum-likelihood method that outputs the overall diversity of a repertoire from measurements on a sample. Recon outputs accurate, robust estimates by any of a vast set of complementary diversity measures, including species richness and entropy, at fractional repertoire coverage. It also outputs error bars and power tables, allowing robust comparisons of diversity between individuals and over time. We apply Recon to *in silico* and experimental immune-repertoire sequencing datasets as proof of principle for measuring diversity in large, complex systems.

## Introduction

Recent technological advances are making it possible to study B- and T-cell repertoires in unprecedented detail^1^. Of special interest is repertoire diversity, defined as the number of different B- or T-cell receptors on cells present in an individual, tissue (e.g., peripheral blood, bone marrow), tumor (e.g., tumor-infiltrating lymphocytes), or cell subset (e.g., influenza-specific IgG^*^ B cells). This interest follows observations that immune-repertoire diversity correlates with successful responses to infection, immune reconstitution following stem-cell transplant, the presence or absence of leukemia, and healthy vs. unhealthy aging^2–5^. The reliability of such observations depends on the ability to measure diversity—and differences in diversity—in overall B- or T-cell populations accurately and with statistical rigor from clinical and experimental samples. Similar requirements also arise in the study of cancer heterogeneity, microbial diversity, and high-throughput sequencing, as well as beyond biology.^6–9^ However, measuring diversity is more complicated than it may seem, for three reasons.

First, “diversity” may refer to any of several different measures. The most familiar diversity measure is the number of different species in a population: the species richness. An example of species richness is the number of B-cell clones in an individual (where “clone” denotes cells with a common B- or T-cell progenitor). Other diversity measures provide complementary information about the size-frequency distribution of species in the population. For example, the Berger-Parker index (BPI) measures clonality, i.e., the dominance of the single largest clone (Fig. 1).^10^ Diversity measures that have been used on immune repertoires include species richness, Shannon entropy (henceforth “entropy”), and the Simpson and Gini-Simpson indices^11–14^. Of these, species richness is unique in that it takes no account of the frequency of each species. In contrast, entropy and other measures systematically down-weight or undercount rarer clones. The above measures (and many more) are related through a mathematical framework described by Hill^15, 16^. Using simple mathematical transformations, this framework allows each measure to be interpreted as the “effective number” of species of a given frequency, facilitating comparisons among different measures (Fig. 1b). For example, entropy, conventionally measured in bits, is converted into an effective number via exponentiation. Thus in the overall repertoire in Fig. 1, the effective number of clones is 7.4 by entropy and 2.9 by BPI (Fig. 1b). The point here is that different diversity measures provide complementary information: two distinct repertoires can have the same species richness but different entropies or BPIs, and vice versa (Fig. 1 d).^10^ Thus, no single measure is likely to capture all of the features of interest in a given repertoire. Consequently, methods for measuring immune-repertoire diversity should be capable of outputting any diversity measure.

**Figure 1.**
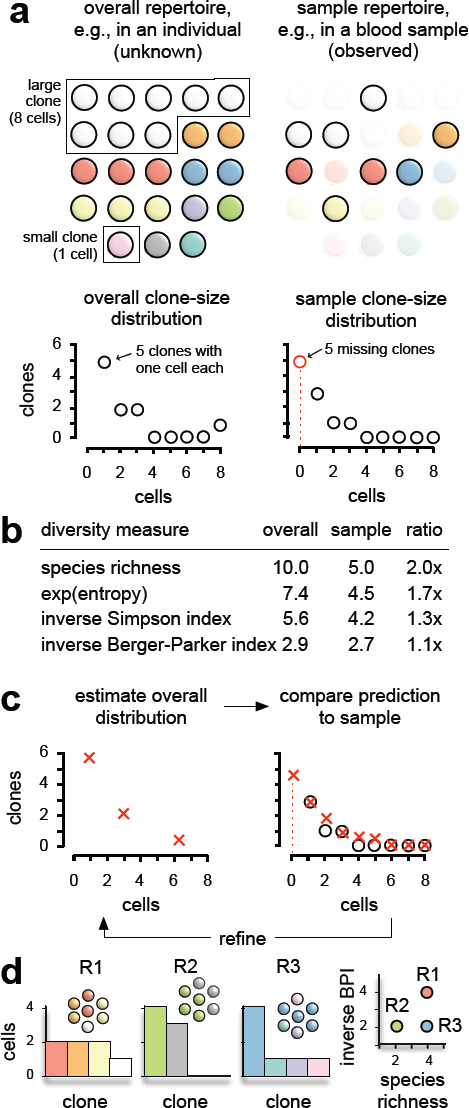
Overall repertoires vs. samples. (a) shows an overall repertoire (top left) and a repertoire from a random sample of this repertoire (top right), together with respective clone-size distributions from the overall repertoire and sample (bottom). Each circle denotes a cell; different colors denote different clones. Note that five clones are missing from the sample entirely, represented by the open red circle at a clone size of zero in the sample clone-size distribution. (b) Sample diversity underrepresents overall diversity across a range of diversity measures. (c) Recon reconstructs the overall repertoire by estimating the number of missing clones and iteratively updating until the predicted clone size distribution in the sample (red crosses) matches the observed clone-size distribution in the sample (open circles), stopping short of overfitting. (d) Different diversity measures are complementary. Repertoires R1, R2 and R3 each have a total of 7 cells. R1 and R3 have the same species richness but different inverse Berger-Parker index (inv. BPI); R2 and R3 have the same Berger-Parker index but different species richness.

Second, the diversity of a sample (e.g. a 5-milliliter clinical blood sample) can differ markedly from the diversity of the overall repertoire from which it derives (e.g., the 5 liters of blood in the body). Although blood and tissue samples may contain thousands or millions of B or T cells, these are only a fraction of the billions of such cells that may comprise an overall repertoire. Consequently, some clones in the overall repertoire, especially small clones, almost always go unsampled and thereby undetected in measurements on samples (Fig. 1a). As a result, sample diversity usually underestimates true diversity (Fig. 1b). This phenomenon is known as the missing-species problem^17^. Weighted diversity measures (e.g., entropy) are less sensitive to missing species than is species richness, since they down-weight the small clones that are most likely to be missing. However, using weighted measures as a substitute for species richness has drawbacks. First, it is unclear what information is lost or biased by selectively ignoring small clones. Second, even using weighted measures, sample diversity will approximate overall diversity only when clone sizes (the number of cells per clone) in the sample approximate clone sizes in the overall population; however, clone sizes will inevitably be biased by the phenomenon of sampling noise. Note that unlike experimental error, which can be minimized, sampling noise is intrinsic to sampling, and will affect measurements even under perfect experimental conditions (e.g. even if every cell in a sample is counted and perfectly annotated). Consequently, depending on the clone-size distribution and diversity measure, sampling can misrepresent overall diversity even when using weighted measures (Fig. 1b and below).

Third, real-world experiments will always exhibit some degree of experimental error, which manifests as noise in sample measurements. Sources include quantitation error due to imprecise cell counts, amplification dropouts, and jackpot effects; sequence error from amplification and sequencing; and annotation error introduced during data processing. Measuring diversity accurately requires methods that address not only the missing species problem and sampling noise, but experimental noise as well.

Existing methods for addressing the missing species problem either output only a single diversity measure (species richness) for the overall population, or else have known or suspected problems scaling to the complexity of immune repertoires. The first category includes Fisher’s gamma-Poisson mixture method, a parametric method that has been used on T-cell repertoires, which involves a divergent sum that can result in large uncertainties^18, 19^; the phenomenological approach of extrapolating from curve fitting^13, 14, 20, 21^; and the Chao estimator (CE), a fast and simple calculation that avoids divergent sums and has been widely used in ecology^22, 23^. The second category includes maximum-likelihood approaches such as the state-of-the-art methods of Norris and Pollock (NP)^24, 25^ and Wang and Lindsay (WL)^26^; however to our knowledge these have not been tested on, or are known not to scale to, highly complex populations like repertoires; or else make restrictive assumptions about the clone-size distribution of the overall repertoire and therefore are not generalizable^27^. Moreover, because a higher-likelihood fit can often be had by adding more small clones, existing maximum-likelihood approaches yield estimates that may overestimate diversity by orders of magnitude or be entirely unbounded-i.e., they may find that best estimate of diversity in the overall population is infinity^28^.

We move beyond these shortcomings using a new algorithm, Recon—<u>r</u>econstruction of <u>e</u>stimated <u>c</u>lones from <u>o</u>bserved <u>n</u>umbers—a generalized high-performance modified maximum-likelihood method that makes no assumptions about clone sizes or clone-size distributions in the overall repertoire, estimates any diversity measure, and leads naturally to sensible error bars that facilitate practical, statistically reliable comparisons between samples, including between individuals and over time, for complex populations.

## Results

### Description

Recon is based on the expectation-maximization (EM) algorithm^6, 29^. Briefly, an initial description of the overall distribution is refined iteratively based on agreement with the sample distribution, adding parameters as needed until no further improvement can be made without overfitting (Fig. 1c). The result is the overall clone-size distribution that, if sampled randomly, is statistically most likely to give rise to the sample distribution subject to the no-overfitting constraint (Fig. S1). The only assumptions Recon makes are that the overall repertoire is large relative to the sample and well mixed.

The input is the observed clone-size distribution in a sample, provided as list of clone sizes and counts. This is easily generated from sequence data by counting clones that have the same number of sequences in the dataset for (at least semiquantitative sequencing. Recon outputs *(i)* the overall clone-size distribution; *(ii)* the diversity of the overall repertoire as measured by species richness, entropy, or any other Hill measure, with error bars; *(iii)* the number of missing species, with error bars; *(iv)* the minimum detected clone size (below); (v) the diversity of the sample repertoire, for comparison to overall diversity; and *(vi)* a resampling of the overall distribution for comparison to the sample and plots thereof. Recon can be run on tumor clones, microbial species, sequence reads, or other populations, including non-biological ones. Recon can also generate tables for power calculations and experimental design.

Recon embodies six improvements over the previous state of the art. First, to avoid dependence on initial conditions or becoming trapped in local maxima, Recon “scans” a number of initial conditions in each iteration of the algorithm. We verified that scanning produces substantially better estimates of overall clone sizes, missing species, and diversity measurements (Fig. S4). Second, Recon optimizes the average of the two best fits in each round (reminiscent of genetic algorithms). Third, it includes a check to prevent overfitting due to sampling noise. Fourth, it makes no assumptions about the overall clone-size distribution, making it widely applicable. Fifth, it improves over previous maximum-likelihood models in avoiding unbounded uncertainties, for example regarding bounds on overall diversity estimates. And sixth, it is substantially faster (Fig. 2b, c).

**Figure 2.**
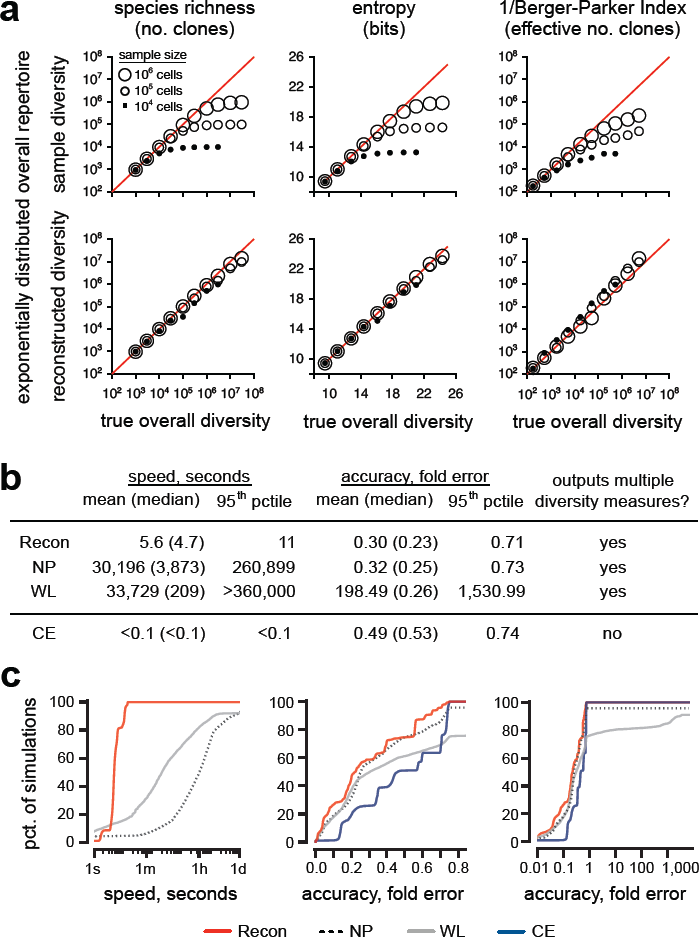
Comparison of diversity estimates. (a) Sample diversity (top) and Recon’s estimate (bottom) of overall diversity vs. true overall diversity for three different sample sizes—10,000 cells (filled circles), 100,000 cells (small open circles), and 1 million cells (large open circles)—for a representative gold-standard distribution without noise (shown in Fig. S2e, left panel; see Fig. S2 for additional examples). Coverage is defined as the number of cells in the sample/effective number of clones in the overall population. Red line, unity (zero error). Left-to-right: species richness, entropy, and the inverse Berger-Parker index. (b) Performance summary of Recon vs. two other state-of-the-art methods for estimating any overall diversity measure (NP and WL) as well as a method for estimating only species richness (CE) on 3,200 noisy distributions, 100 realizations of noise for each of 32 combinations of exponential and multimodal distributions (Online Methods), coverage (0.05-0.3x), and overall diversity (100,000-3 million clones in the overall population). (c) Cumulative distribution of performance for distributions in (b) showing Recon is much faster than NP and more accurate than WL, which could be off by orders of magnitude. (d) Cumulative distribution of performance for distributions in (b) showing Recon is more accurate than CE.

Current methods tend to overestimate species richness when coverage is low, as small clones added to the estimate result in overfitting of the sample distribution—in the limit, as mentioned, leading to an estimate with infinite infinitesimal clones. Recon uses discrete clone sizes, which in the worst case ensures that estimates are bounded by the number of cells in the overall repertoire (clones cannot outnumber cells). Beyond that, Recon’s use of both a noise threshold and the (corrected) Akaike information criterion provide tighter bounds, rejecting additional clones unless their expected contribution to the sample rises above sampling noise (by 3 standard deviations in our implementation) and outweighs the penalty of adding more parameters. The trade-off is that for each sample, there is a minimum clone size that Recon can detect: if ≤ 1, Recon’s species-richness estimate will include clones represented by just a single cell in the overall repertoire, if there are any; if > 1, in principle there may be clones in the overall repertoire that are too small to detect. In this case Recon can be used to calculate a strict upper bound, *U*, on species richness that includes clones that may be “hiding” (Online Methods and Supplementary Information). However we note that even in this case, in practice, for a given sample, the smallest clones detected may still be the smallest clones there are (the case for our *in silico* repertoires; below).

### Validation

We validated Recon on *in silico* repertoires that spanned nearly five orders of magnitude of overall diversity (300-10 million clones) and a wide range of clone-size distributions: from steep, i.e., dominated by small clones, to flat exponentials; reciprocal-exponential distributions that derive from a generative model; and multiple bimodal distributions of small and large clones, 1,711 in all, with and without simulated experimental noise (Online Methods). These repertoires served as gold standards. We sampled a known number of cells from each, for coverage ranging from 0.01x to 10x, and used Recon to reconstruct overall repertoires from each sample. (Coverage is the number of cells in the sample the number of clones in the overall repertoire.) We then compared the diversity of the reconstructed overall repertoire with the true overall diversity and sample diversity. We measured diversity by species richness, entropy, Simpson Index, and BPI (Fig. 1b).

First, to illustrate the extent of the problem Recon solves, we compared sample diversity with overall diversity (Fig. 2a). For a given sample size, higher overall diversity means lower clonal coverage (the number of cells in the sample per clone in the overall repertoire). For each repertoire, the error, defined as the difference between sample and overall diversity, grew as coverage fell below 1x, because samples cannot have more clones than cells. Consequently, for species richness, sample diversity underestimated true diversity by 50% at 1x coverage, 10 fold at 0.1x coverage and 30 fold at 0.03x coverage. The weighted measures performed little better, even for the flattest clone-size distributions that we tested, partly due to the absence of clones large enough to dominate these repertoires (e.g., leukemic clones; Figs. 2 and S2). We concluded that sample diversity is generally an unreliable proxy for true diversity below 1x coverage in the absence of dominant clones.

In contrast, Recon’s estimates of overall diversity showed excellent agreement with true diversity across the range of diversity measures, even at <1x coverage (Fig. 2a, lower panels). For species richness, Recon’s estimates were accurate to within 1% of the true diversity at 10x coverage, 10% at 3x coverage, and 50% at just 0.03x coverage—at which there is just one cell in the sample for every 30 clones in the overall repertoire. Error for entropy and other weighted measures was lower. Recon was also robust to noise (Fig. 2b–c).

To visualize self-consistency, we resampled from the overall repertoires we reconstructed from our gold-standard distributions in order to compare the resulting sample clone-size distributions to those of the original samples. We found excellent agreement between predicted and observed frequencies of clone sizes across the range of overall diversities and levels coverage, including on numbers of missing clones (Fig. 3). Recon’s ability to estimate the number of missing clones accurately was a key contributor to the accuracy of its overall diversity estimates. The number of missing clones depended strongly on the number of singlets (clones represented by a single cell) and doublets (two cells) in the sample: large singlet-to-doublet ratios, with enough of both for low sampling noise, gave more accurate estimates.

**Figure 3.**
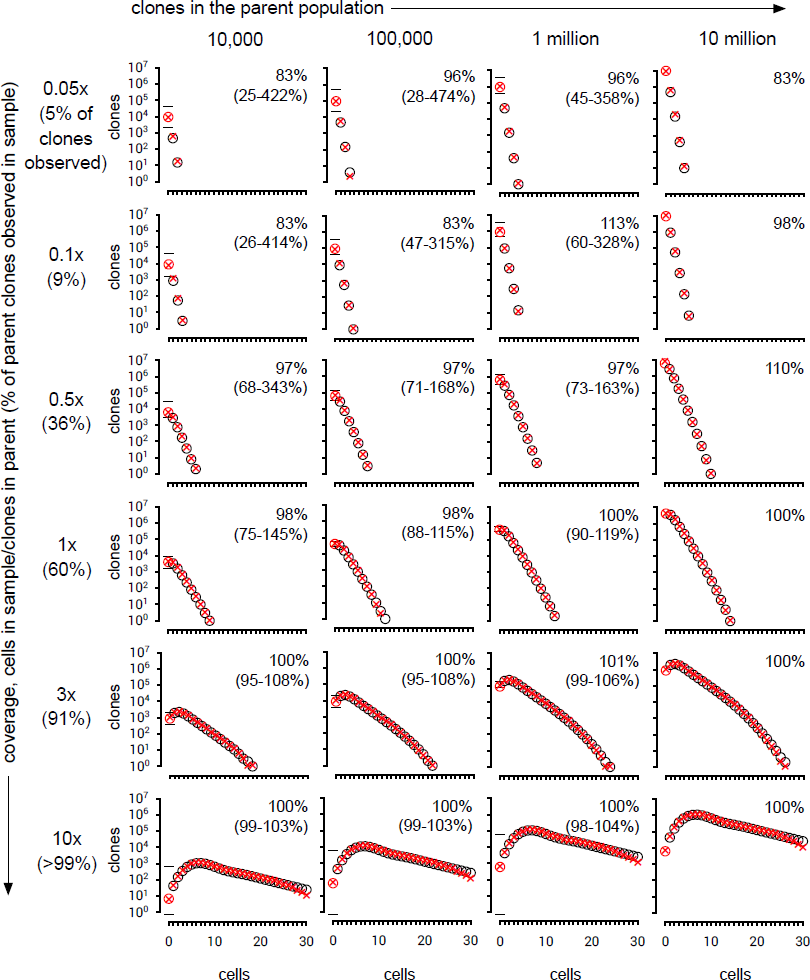
Predictions vs. simulated observations, *in silico* gold standards. Shown are fits to observations from representative gold-standard distributions of the shape shown in Figure S2e, left panel. Left-to-right: overall distributions with increasing numbers of clones. Top-to-bottom: increasing sample size measured in coverage of the number of clones in the overall population. Open black circles denote observed clone-size distributions, which was the input data given to Recon. The open red circle denotes the number of missing clones, which was not known to Recon. Red crosses denote Recon’s prediction of the clone-size distribution in the sample, based on its reconstruction of the clone-size distribution of the overall repertoire. This includes a prediction for the number of missing clones, plotted as the number of clones of size zero, with error bars as shown.

In head-to-head comparisons on 3,200 *in silico* samples with experimental noise (Online Methods), Recon was both faster and more accurate than the prior methods NP and WL, which like Recon can be used to estimate overall diversity by multiple diversity measures (Fig. 2b–c; Fig. S2h). Specifically, Recon’s median runtime of 4.7 seconds (95^th^ percentile, 11 seconds) was >40x faster than WL and >800x faster than NP, both of which often took hours and sometimes days to complete (Fig. 2c). Recon’s median error of 0.23x was smaller than that of NP (0.25x) and WL (0.26x), which was often off by orders of magnitude (mean, 198x; 95^th^ percentile, >1,500x). Recon was also more than twice as accurate as CE (0.53x median error; Fig. 2b, e), which is fast but limited to outputting species richness.

### Error bars and power calculations

Detecting reliable differences in overall diversity requires that diversity estimates have reliable bounds. Recon outputs two types of bounds: error bars on overall diversity (more precisely, on the effective number of clones greater than or equal to a minimum detected clone size) and a maximum-possible overall species richness, *U* (Supplementary Information).

To build error bars, we first sampled gold-standard repertoires systematically across three orders of magnitude of coverage (0.01x-10x). For each sample, we used Recon to estimate overall diversity. Because higher coverages produce better estimates, the resulting error profile converges with increasing coverage to the true overall diversity (Fig. 4a). The upper and lower contours of this profile correspond to the largest and smallest values of estimated diversity that are consistent with a given true diversity. To make an error bar for a given estimated diversity, Recon uses the contours of the error profile to find the true diversities for which the estimated diversity is at the lower bound and the upper bound. These respectively define the upper and lower error bars (Figs. 4b, 4c). Following cross-validation, we adjusted our error profile slightly so that error bars reflect 95% confidence intervals (Online Methods). Combining error profiles across all samples suggests that ≥1x coverage generally produces error bars of ≤10% for overall species richness (Fig. 4d), consistent with our previous observations (Fig. 2).

**Figure 4.**
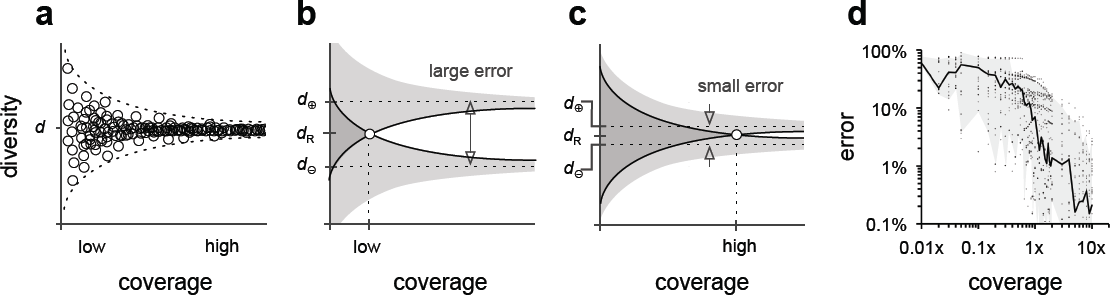
Error bars. (a) shows a schematic representation of Recon’s diversity estimates (open circles) from a single gold-standard *in silico* repertoire with overall diversity *d* for many different levels of coverage (=sample size/d). We used the absolute value of the proportional error of the worst fit at each level of coverage, making an error profile that is vertically symmetric around *d*. Given a test sample, Recon first estimates the overall diversity, d_R_, and the coverage. (b) Using the error profile, it then looks up the maximum (d_e_) and minimum (d_e_) diversities that are consistent with its estimate (d_R_); schematically, this is where the edges of the funnel plots for d_e_ and *de* intersect. (c) Higher coverage gives smaller error (arrows). (d) Combining errors from all 1,711 gold-standard repertoires into a single plot suggests that >1x coverage generally gives error bars of 5-10% for species richness (line, median; shaded area, 5^th^-95^th^ percentiles).

Recon uses this error-bar framework to determine the coverage required to confidently detect differences in diversity between samples (e.g., between individuals or over time). Given an order-of-magnitude estimate of the overall diversity for two samples, it outputs the minimum sample size for which error bars for overall diversity estimates from these samples would not overlap, at detection thresholds ranging from e.g. 1.1x to 5x (Table 1). This sample size is the minimum required to reject the null hypothesis that two estimates that differ by a given amount are actually from the same overall repertoire, at a confidence level of p = 0.05 (Supplementary Information). Not surprisingly, detecting larger differences requires smaller sample sizes; less obviously, for a given overall diversity there is a minimum sample size below which the number of non-singlets is expected to be too small for Recon to run. So an experiment designed to detect a 1.1x (10%) difference in species richness between two samples, in which the samples are drawn from overall repertoires that have ∼100,000 clones, will require ≥313,792 cells from each sample for analysis. This is the number of cells in the sample that are in small (≤30 cell-) clones that Recon requires to perform reconstruction; if half of the cells in a sample of 314,000 cells belong to a single large clone, e.g. because of leukemia, the remaining half comprising the non-leukemic clones will be sufficient to detect a 20% difference in the species richness of the non-leukemic portion of the repertoire (which requires ≥153,543 cells), but not 10%.

**Table 1.** Power calculations. Table entries give the minimum number of cells that must be analyzed in order to be able to detect a given fold-difference in species richness between two samples at p=0.05 (row headings), given an expected overall species richness (column headings). As noted in the main text, these numbers exclude cells that might belong to large clones (here, of clone size ≥30 in the sample). Minima required for reliable reconstructions are in gray. See Supplementary Information for details.

|  | 10,000 | 30,000 | 100,000 | 1 million | 3 million |
| --- | --- | --- | --- | --- | --- |
| 1.1 | 34,634 | 118,418 | 313,792 | 2,211,303 | 16,230,339 |
| 1.2 | 19,103 | 56,989 | 153,543 | 1,277,637 | 10,598,339 |
| 1.3 | 14,142 | 28,206 | 85,156 | 649,124 | 1,947,385 |
| 1.4 | 14,142 | 27,711 | 70,415 | 639,665 | 1,919,012 |
| 1.5 | 14,142 | 27,238 | 64,982 | 630,590 | 1,891,799 |
| 2.0 | 14,142 | 24,495 | 64,977 | 510,381 | 1,524,687 |
| 5.0 | 14,142 | 24,495 | 44,721 | 141,421 | 244,949 |

To test Recon and our error-bar framework beyond exponentially and multimodally distributed samples, we ran it on a sample distribution previously identified as causing difficulties for overall species-richness estimation by multiple existing methods, corresponding to an overall population of ∼3,000 species sampled at ∼0.8x coverage (Supplementary Information)^28^. Three- and four-point mixture models, a logit normal model, a log-gamma model, and a beta model gave variable estimates that ranged from 2,930-3,494 overall species, with non-overlapping error bars that ranged from 2,867 to >10,000. In contrast, Recon returned an estimate of 3,014 overall species, with error bars (2,709-3,513) that bracketed the range of other models’ estimates, suggesting Recon improves on multiple methods beyond WL and NP in arbitrary and/or difficult cases.

### Experimental data

Having validated Recon, we next applied it to six experimental datasets: four of paired heavy-and-light chain sequence and two of heavy chain sequence (Online Methods). We used the authors’ clone definitions—clusters of reads with ≥96% nucleotide identity in heavy-chain complementarity determining region 3 (CDRH3)^30^ or reads with identical CDRH3s and V_H_ annotations ^31^—with the caveats that clone assignment is difficult, some cells may not have been sequenced, artifacts are possible, and sequencing is only semi-quantitative. Because such datasets reflect the current state of the art in the field and are used for diversity measurements, we considered them as (imperfect) samples and used Recon to estimate diversity for the corresponding overall repertoires (Table 2). As with our gold-standard samples, resampling showed excellent agreement with the observed data (Fig. 5). For four of the six repertoires, we found that missing species accounted for the majority of clones: i.e., half of all clones are unseen, and species richness in the sample underrepresents overall species richness by 2x. Entropy was generally very similar between samples and overall repertoires, resulting from very large clones and/or PCR jackpot effects that contribute disproportionately to the entropy calculation. Thus in these datasets, overall species richness, estimated using Recon, captures information lost during sampling that entropy does not.

**Figure 5.**
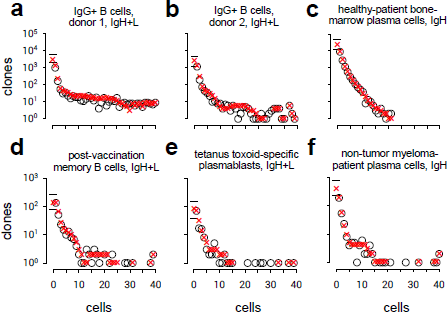
Predictions vs. observations, experimental data. Shown are Recon’s estimates of overall diversity for six experimental datasets. These included (a, b) immunoglobulin heavy (IgH)- and light-chain (IgL) paired-chain sequencing experiments from IgG^*^ B cells from the blood of two different subjects, (c) pooled-DNA IgH sequencing experiments on the bone-marrow plasma cells from a healthy adult, (d) IgH^*^L of post-vaccination memory B cells, (e) IgH^*^L tetanus toxoid-specific plasmablasts, and (f) pooled-DNA IgH sequencing experiments on the bone-marrow plasma cells from a multiple myeloma patient (only the non-myeloma cells). Details, including references, are presented in Table 2.

**Table 2.** Diversity estimates for experimental datasets from humans. Summarized are Recon’s estimates of overall diversity for six datasets; its estimate of the number of missing species; comparisons to sample diversity, for species richness and entropy (given as effective numbers; 2^bits^); the minimum detected clone size (see main text); and upper bound for species richness that includes potential “hiding” clones. Cell-surface phenotypes were as follows: IgG^*^ B cells, IgG^*^CD2^−^CD14^−^CD16^−^CD36^−^CD43^−^CD235a^−^; post-vaccination memory B cells, CD19^*^CD3^−^CD27^*^CD38^int^; tetanus-specific plasmablasts, CD19^*^CD3^−^CD14^−^CD38^**^CD27^**^CD20_; plasma cells, CD138^*^. See references for details.

| Subset | Source | Method | Cells | Species richness |  | Missing species | Entropy (eff. no.) |  | Min clone size, cells | Strict upper bound, clones |
| --- | --- | --- | --- | --- | --- | --- | --- | --- | --- | --- |
|  |  |  |  | Sample | Overall |  | Sample | Overall |  |  |
| IgG<sup>+</sup> B cells, individual 1<sup>22</sup> | healthy adult | IgH+L single-cell | 61,000 | 2,759 | 5,870 (4,761-8,395) | 3,111 (2,002-5,636) | 696 | 700 (691-720) | 400 | 1 million |
| IgG<sup>+</sup> B cells, individual 2<sup>22</sup> | healthy adult | IgH+L single-cell | 47,000 | 2,211 | 4,616 (3,374-7,000) | 2,405 (1,163-4,789) | 345 | 348 (327-373) | 700 | 5 million |
| memory B cells (IgG, IgM, and IgA)<sup>22</sup> | healthy adult vaccinee | IgH+L single-cell | 8,000 | 336 | 473 (446-614) | 137 (77-245) | 21 | 21 | 30,000 | 14 million |
| tetanus toxoid-specific plasmablasts<sup>22</sup> | healthy immunized adult | IgH+L single-cell | 2,000 | 159 | 239 (200-313) | 80 (41-154) | 3.5 | 3.5 | 1,000 | 300,000 |
| bone-marrow plasma cells<sup>24</sup> | healthy adult | IgH pooled DNA | 26,000 | 14,337 | 37,110 (27,350-58,916) | 22,773 (13,013-44,579) | 11,148 | 21,582 (20,891-22,572) | 80 | 4 million |
| non-tumor plasma cells<sup>24</sup> | multiple myeloma patient | IgH pooled DNA | 30,000 | 325 | 703 (563-1,081) | 378 (238-756) | 1.4 | 1.4 | 80 | 80,000 |

## Discussion

High-throughput technologies enable highly detailed descriptions of B- and T-cell repertoires. That these descriptions are generally of samples, and not e.g. blood or tissue repertoires overall, may seem to be a distinction without a difference when samples contain many cells. However, and perhaps counterintuitively, it turns out to be critical for estimating overall diversity. Unless the number of cells in a sample exceeds the number of clones in the overall repertoire by ∼3-10-fold (Fig. 3), sample and overall diversity may bear little relation (Figs. 2a, S2a–c). Importantly, this discrepancy is not a technological shortcoming but an inherent constraint of random sampling (Fig. 1a). In humans, overall repertoires may contain many millions of clones. Because routine blood samples rarely contain more than a few million B and T cells of any sort combined, they are too small for sample diversity to serve as a reliable proxy for overall diversity. Thus conclusions drawn only from sample diversity measurements warrant caution.

This caveat applies for all diversity measures. Entropy, often used to measure sample diversity in immune-repertoire studies, is less prone to undercounting. However, in our gold-standard repertoires even BPI, the Hill measure least prone to undercounting and most robust to missing species, underestimates overall diversity by an order of magnitude for levels of coverage encountered in experiments (Figs. 2, S2). It is unsurprising, then, that sample entropy can also underestimate overall entropy in these repertoires (Figs. 2, S2). Additional caveats apply to experimental datasets. Insufficient read clustering will overestimate species richness; for clone sizes defined proportional to the number of reads, PCR jackpot effects can produce artificially large “clones,” overestimating entropy. These biases, not mutually exclusive, may affect species richness and entropy in the experimental datasets we studied (Table 2). Better quantitation (e.g., via barcoding and robust clonality modeling) would mitigate these biases but not the bias intrinsic to sampling, which Recon addresses.

Recon outperforms prior methods even for large, complex clone-size distributions, at fractional coverage, and in the presence of experimental noise (Figs. 2-3, S2). Notably, Recon avoids WL’s major failure modes: the 10-50 percent of cases in which WL unpredictably takes hours or days to run and/or overestimates diversity by orders of magnitude. Recon’s characteristic runtime of seconds to a minute is especially faster than NP, and negligible relative to the hours-to-days of current sequence-processing pipelines. These advantages are not unexpected given that Recon was designed for handling samples from large, complex, and arbitrary distributions.

Error bars and power tables are necessary steps toward being able to compare diversity between samples and over time and thus for evaluating diversity as a potential biomarker. Recon’s error bars and tables for entropy, BPI, and other measures mean differences can be assessed for any measure or noise level. Recon’s error bars perform well by practical tests, bracketing the number of missing species in validation studies and squaring previous models^28^. Its power tables offer guidance for sample requirements during experimental design and suggest expected limitations for different studies. For example, measuring the species richness of naive repertoires of ∼10^7^ clones^31, 32^ will likely require phlebotomy or apheresis samples; even then, detecting 50-percent differences is probably the limit (Table 1). Meanwhile, measuring diversity for effector/memory subsets should require only routine blood draws (2-6mL), which should detect sub-fold differences. For marrow, spleen, tumor, granuloma, or abscess samples, the investigator must decide whether the sample is well mixed, which Recon requires.

High-throughput technologies hold much promise for measuring diversity in repertoires, cancer, and other complex populations, but current limitations warrant caution. Because most sequencing experiments are still only semi-quantitative, the number of reads does not always reflect the number of cells. Chimerism and sequencing/annotation errors mean not all clusters are clones. Incomplete cell lysis and sequencing inefficiencies can underestimate sample size. These limitations affect the calculation and interpretation of diversity estimates and upper bounds; the examples we have shown should be interpreted accordingly, even as they illustrate application of our method. Overcoming these limitations will improve our understanding of overall diversity, a defining characteristic of complex systems that we can now better measure.

## Online Methods

### Core algorithm

Mathematically, the problem is to find the B- or T-cell clone-size distribution in the individual (the “parent” or “overall” distribution) that is most likely to give rise to the clone size distribution that is observed in the sample (the sample distribution) (Fig. 1). From the parent distribution, we can then calculate overall diversity according to any diversity measure in the Hill framework. The core of our method is the expectation-maximization (EM) algorithm, in which a rough approximation of the parent distribution is refined iteratively until no further improvement can be made without overfitting^29^.

The EM algorithm begins by assuming a parent distribution in which clones are all the same size, taken from the mean of the observations. To perform the fit, we need to know not just the observed clone frequencies but also the number of missing species, which is unknown and therefore must first be estimated. Following previous work^33^, we estimate the number of missing species by calculating the expected clone size distribution for a (Poisson) sample of the parent distribution (see “Sampling” below) and applying the Horvitz-Thomson estimator^34^. We then fit the clone size of the parent distribution using maximum likelihood, recalculate the number of missing species, and repeat these steps until a self-consistent number of missing species is obtained. This completes the first iteration of the algorithm, yielding the uniform parent distribution that is most likely to give rise to the sample distribution.

In the second iteration, we refine this uniform parent distribution by adding a second clone size. We estimate the number of missing species for this new two-size distribution, fit the two clone sizes and their relative frequencies by maximum likelihood, and, as in the first iteration of the algorithm, repeat until there is no further improvement^33^. The result (pending a check for overfitting, below) is the two-clone-size parent distribution that is most likely to give rise to the sample distribution.

In subsequent iterations, we continue to refine the parent distribution by adding clone sizes and refitting as above, iterating until no more clone sizes can be added without overfitting (using the corrected Akaike information criterion [AICc] as a stop condition). The result is the desired MLE. Note that whereas the sample distribution generally traces out a smooth curve, the MLE parent distribution is spiky, reflecting the limited resolution that information in the sample distribution provides about the parent distribution.

### Sampling

We assume that each clone in the individual contributes cells to the sampled population according to a Poisson distribution. This will be true if (*i*) clones are well mixed in the blood or evenly distributed in the tissue being sampled, (*ii*) the parent population is sufficiently large that the Poisson estimate for the probability of e.g. a singleton contributing >1 cell is negligible, and (*iii*) no single clone is a large fraction (∼30% or more) of the parent population. In practice, condition (*iii*) is satisfied by counting large clones directly (see “Fitting”).

### Fitting

The largest clones may be represented by hundreds or even thousands of cells in a sample. For such large clones, sampling error is small: the relative size of the clone in the sample and in the individual will be about the same. As a result, clones that are large enough to have sufficiently small sampling error do not have to be fit by EM, and instead can simply be added to the MLE. We found that using a threshold of 30 cells, and therefore applying EM only to clones that contribute ≤30 cells to the sample and then adding larger clones back to the resulting MLE gives results that are indistinguishable from applying EM on the entire sample distribution, but with vast gains in speed. (Note that not seeing clones of sizes similar to sizes for which clones are seen is itself an observation, and therefore counts toward the number of observations used for calculating the AICc.)

### Scanning

In the standard EM algorithm, the exact sizes and frequencies of clones in the final MLE can vary depending on the sizes and frequencies used at the start of each iteration, reflecting different relative maxima. To find global maxima, we developed a “scanning” approach in which we applied EM to many starting clone sizes and frequencies (56 in our implementation), ranking results by maximum likelihood (after first adjusting likelihoods according to the number of ways to choose clones in each distribution; see Supplementary Information). In each round we perform an additional fit with starting clone sizes and frequencies at an average of the two top-ranked results. We then select the resulting best-ranked fit from the starting points. Runtime and (to some extent) accuracy correlate with the number of starting points.

### Diversity measures

Species richness, entropy, the Gini-Simpson Index, BPI, and indeed many other diversity measures are related to each other through the mathematical framework of the so-called Hill numbers^15, 35^. These form a series in which the index reflects the extent to which counts are weighted toward large clones. Species richness, in which large and small clones are counted equally and so large clones are unweighted, has an index of zero and is denoted ^0^*D* (“D-zero”). Other measures, or simple mathematical transformations thereof, correspond to larger indices; these include entropy (ln(^1^*D*)), the Simpson Index (1/^2^*D*), and BPI (1/^∞^*D*).

We calculated ^0^*D*, ^1^*D*, ^2^*D*, and ^∞^*D* for sample and overall distributions from *in silico*-sampled synthetic gold-standard distributions (see “Validation” below and in the main text) and from several published data sources (see “Experimental Data” in the main text). These *^q^D* are a function of frequencies of clone frequencies *p_i_*, where *i* ranges over each clones and the frequencies are normalized to 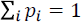, defined as 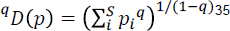

We calculated ^0^*D* by simply counting the number of different clones, ^1^*D* according to 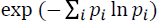, ^2^*D* according to the definition, and ^∞^*D* as the reciprocal of the frequency of the largest clone (the above definition reduces to these expressions for the value *q* = 0 and in the limits *q* → 1 and *q* → ∞).

### Validation

We validated Recon against CE, Norris and Pollock, and Wang and Lindsay by generating a wide range of biologically plausible synthetic parent distributions of 10^9^ cells *in silico*, sampling from these distributions to produce samples of different known sizes, using the samples to estimate overall diversities according to species richness by the listed methods and the other above measures for all but CE (which outputs only species richness), and comparing these estimates against the (known) calculated diversities of the original parent distributions. We studied three families of test distributions in detail: (*i*) exponential distributions (of the form 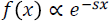, where *x* denotes clone size, *f*(*x*) is the frequency of clones of that size, and *s* is a parameter that controls the steepness of the distribution), which are simple distributions that describe the shape of observed sample distributions phenomenologically; (*ii*) “reciprocal-exponential” distributions 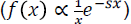, which are the analytical solution to a simple biologically plausible model of the dynamics of most B- and T-cell clones; (*iii*) bimodal distributions with the largest clones an average multiple of the size of the smallest clones (e.g. 20-30x) in the overall population. We tested these distributions systematically by varying the steepness from very steep (*s*=1.2) to nearly flat (*s*=0.12) exponential distributions and different multiples for the bimodal distributions, encompassing the a range of biologically plausible clone-size distributions, with and without noise. We investigated three different modes of noise: (*i*) noise added to each count *n* with mean of zero and standard deviation 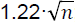; (*ii*) a small baseline amount of noise added to all clone sizes; and (*iii*) sporadic noise at random clone sizes (reminiscent of PCR jackpot effects). For completeness, we tested on both Macintosh (2.7GHz Intel Core i5 running OS X 10.11.1) and Linux (2.3-2.8GHz Intel Xeons running RHEL CentOS 6.6) platforms. NP and WL fits that were still incomplete after 100 hours were terminated.

### Error bars

Error bars define the range of overall diversity values that, given inevitable sampling error and any error in reconstructing parent distributions from samples of a given size, are consistent with Recon’s estimate. We determined error bars for each diversity measure (species richness, entropy, etc.) as follows (Fig. 4). First, we generated a wide range of exponentially and multimodally distributed *in silico* parent populations with known diversities of 3×10^2^-1×10^7^ species. Next, we took samples of these known distributions at systematically increasing coverage/sample sizes from 0.01× to 10× and, for each sample size, ran Recon to estimate the overall diversity, running on 1,716 samples in all (Fig. 4a). Five outliers (0.3%) were removed, leaving 1,711. For each overall diversity and coverage, the error was defined as the difference between the (true) overall diversity and Recon’s overall diversity estimate. Given a test sample, the coverage, and Recon’s estimate, one can then look up or interpolate from these errors the largest and smallest diversity values that are consistent with the estimate (Fig. 4b, c). These upper and lower bounds define the desired error bars on Recon’s estimate.

We established these error bars as 95% confidence intervals using Monte Carlo cross validation. Briefly, we randomly partitioned the above 1,711 samples 70-30 into reference and validation sets 100 times, each time using the reference set to calculate error bars for the samples in the validation set and counting how often error bars bracketed true diversity. These raw error bars bracketed true diversity in 93.6±1.3% of cases; adjusting them by raising the upper bar by 1.6 percent brought this figure to the desired conventional level for confidence intervals, 95% (96.2±1.0%). Note that error bars bracketed true diversities despite the formal possibility of there being clones in the parent population too small to observe in the sample (see Minimum detected clone sizes and upper bounds (*U*), below), meaning in practice this was not an issue. The above procedure can be generalized to incorporate arbitrary models of experimental noise.

### Experimental datasets

We found and downloaded six publically available datasets. Four were from paired heavy-and-light-chain sequencing experiments: two of IgG^*^B cells (from two subjects), one of memory B cells post-influenza vaccination, and one of tetanus-toxoid-specific plasmablasts^30^. Following that study’s methods, we clustered reads with ≥95% heavy-chain complementarity-determining region 3 (CDR3) nucleotide identity (the study treated clusters as clones). The other two datasets were of pooled PCR of heavy-chain genomic DNA from bone-marrow plasma cells from a healthy subject and non-myeloma plasma cells from a subject with multiple myeloma, with clones defined as sequences with identical CDR3s at the amino acid level and identical V_H_ nucleotides^36^. We estimated the total number of IgG^*^ B cells, post-vaccination memory B cells, tetanus-specific plasmablasts (and plasma cells), bone-marrow plasma cells in a healthy patient, and non-myelomatous plasma cells to be 75 million, 260 million, 3.5 million, 6 million, and 3 million, respectively, for *N* (See below)^37–42^.

### Minimum detected clone sizes and upper bounds (U)

The smallest clone size in the reconstructed clone-size distribution is described by two parameters: the mean number of cells that each clone of this size contributes to the sample, *m*_min_, and the fraction of all clones that are of this size, *w*_m_. The size of this smallest detectable clone in the overall repertoire is *m*_min_ scaled to the total number of cells: *m*_min_*N/S*. This is Recon’s minimum detected clone size. It is possible that there are clones smaller than this size in the overall repertoire, but because they contribute a mean of zero cells to the sample they are not detected and therefore do not contribute to Recon’s estimate of overall species richness. An upper bound on species richness that includes clones smaller than the minimum detected clone size, *U*, is obtained by assuming that all cells in clones that could be smaller than this are singlets: *U* = *R*_max_W_m_m_min_*N/S*, where *R*_max_ is Recon’s upper error bar estimate of overall species richness (Supplementary Information). We calculated these quantities for our validation and experimental data.

## Acknowledgements

The authors thank the reviewers, William A. Link for correspondence, and Rima Arnaout for critical reading of the manuscript. This work was supported by the National Institutes of Health (NIH) National Institute of Allergy and Infectious Disease (NIAID) grant K08AI114958-01 (RA), Ruth Moorman and Sheldon Simon (RA), and American Heart Association (AHA) grant 15GPSPG23830004 (RA).

---

**Figure S1.** The Recon algorithm. Steps in the flowchart are as described in the main text, Online Methods, and Supplementary Information.

**Figure S2.** Recon diversity vs. other estimates showing fits to additional gold standard repertoires plotted as for Figure 2. (a)-(c) Comparisons of sample diversity (top) to Recon diversity (bottom) plotted as in Figure 2a for (a) a steep exponential clone size distribution (b) a bimodal distribution in which the overall distribution contains a population of small clones and a population 31 times as large and (c) a bimodal distribution in which the overall distribution contains a population of small clones and a population 20 times as large. (d)-(g) Comparison of species richness estimates by Recon (middle) and CE (right) shown as in Figure 2b for an example additional gold standard overall distributions (left) for (d) a steep exponential clone-size distribution, (e) a shallow exponential clone-size distribution, (f) a bimodal distribution in which the overall distribution contains a population of small clones and a population 31 times as large, and (g) a bimodal distribution in which the overall distribution contains a population of small clones and a population 20 times as large. (h)-(k) Comparative performance of Recon, NP, WL, and CE for noisy samples from steep exponential distribution shown in (d) at 0.05x for overall populations with (h) 10 million, (i) 3 million, (j) 100,000, and (k) 10,000 clones. (l-m) Comparative performance for higher D numbers on (l) the exponential distribution in (d) at 0.05x coverage and (m) the bimodal distribution in (f) at 0.3x coverage. (Note CE outputs only *^0^D*, species richness.)

**Figure S3.** Recon vs. CE, NP, and WL on noisy distributions. Each pair of cumulative distributions show accuracy (left) and speed (right) for 100 realizations of noise on the different distribution types described in Fig. 2.

**Figure S4.** Scanning. Probability densities of the ratio of estimated missing species/true missing species demonstrating the benefit of using additional starting points. Fits using, in each round of fitting, 9 (red), 20 (yellow), 56 (green), 72 (pink) and 110 (blue) combinations of starting weights and means (yellow) show that the set of 56 starting points used in the main study result in a sharper peak of the probability distribution function (pdf) near 1.0, and diminished trapping in local minima away from 1.0. Pdfs are plotted using Gaussian kernel density estimates over 800 samples from gold-standard distributions (see main text).

